# Genetic Convergence Analysis of CRISPR Perturbations Deciphers Gene Functional Similarity

**DOI:** 10.1101/2025.11.13.688060

**Authors:** Tianyu Zhang, Ergan Shang, Kathryn Roeder

## Abstract

Pooled CRISPR screens with single-cell RNA sequencing readout (Perturb-seq) have emerged as a key technique to determine the functionality of a gene by directly perturbing the DNA of the gene. One of the most intriguing recent problems is quantifying the similarity between CRISPR perturbations, for example, whether they upregulate the same set of downstream genes. In this context, genetic convergence refers to the phenomenon where CRISPR disruptions of different genes lead to a similar downstream outcome. Existing methods are mostly heuristic. We present XConTest, a two-step, cross-validated procedure for assessing the genetic convergence problem. The test statistics calculated from that procedure are approximately standard normal when the two perturbations have an orthogonal influence on the cell expression profile. We apply XConTest to two studies: an investigation of the common impact of a suite of autism genes, and a large-scale study of genes associated with immune response to determine sets of genes with common functionality.

## Introduction

Recent advances in single-cell transcriptomics, coupled with high-throughput CRISPR-based perturbation platforms, including Perturb-seq [1, 2] and CROP-seq [3, 4], have enabled systematic functional interrogation of genes at scale. These methodologies facilitate direct assessment of gene function by measuring transcriptional responses to targeted gene knockouts or knockdowns at single-cell resolution. Analysis of these responses permits grouping perturbed genes into co-functional clusters, inferred from the similarity (or *convergence*) of their downstream effects [2, 5, 6, 7, 8, 9, 10]. Such clusters often reflect coordinated regulation of gene expression modules, revealing the architecture of gene networks underlying cellular phenotypes.

Despite these technological advances, both conceptual and methodological gaps remain in quantifying genetic convergence. Convergence reflects the modular organization of cellular regulation—distinct perturbations may impact on the same signaling cascade or regulatory subnetwork. Conceptually, this similarity extends beyond effects of the same direction: perturbations that induce opposing transcriptional changes within a shared module may still operate through a common regulatory axis [11, 12, 13]. The relevant signal lies in the alignment of affected pathways rather than the mere concordance of mean shifts, pushing the definition of convergence beyond simple similarity in average expression changes. Methodologically, detecting such convergence requires isolating structured perturbational effects from background variability. A scalable statistical framework is particularly relevant when both the number of perturbations and the transcriptomic dimension are large.

Most available strategies for identifying genetic convergence build on a common fundamental principle: sets of perturbations tend to cluster in their effect on clusters of down-stream genes, i.e., biclustering. The downstream genes are typically pre-clustered based on correlated gene expression [9] or correlated perturbation effect size [8]; these sets are called gene modules or gene programs. Next, perturbations with similar effects on modules are identified, either by clustering effect sizes [6, 8] or by identifying those perturbations with a significant impact on pre-specified modules [9, 10]. Another approach developed in [7] and [14], computes distances between perturbations—or directly examines the average expression per gene after perturbation—to cluster similar perturbations directly. This approach does not emphasize contrasting the perturbations to the control, and hence lacks the power to detect differences/similarities in perturbations. The signal is swamped by the noise of enormous baseline genes that are not impacted by the perturbation. However, none of the available methods provides a measure of significance for convergence itself.

Here, we introduce XConTest, a framework for discovering and validating genetic convergence. XConTest allocates a subset of cells to train linear classifiers that capture shared differential expression (DE) patterns (identical or reversed directions), while reserving held-out data to ensure discovery is replicable. A pair of perturbations is determined to be convergent when (1) there exists a set of genes whose expression levels are consistently modified in both perturbations against the non-targeting and (2) the trained classifier can effectively discriminate the perturbed cell from the control in the held-out independent samples. A cross-validation–based design further maximizes statistical efficiency by aggregating across multiple training–validation splits. Through simulations, we examined the statistical properties of XConTest and demonstrated its utility in two Perturb-seq datasets. We also compare XConTest with another novel method that is naturally derived from current DE analysis (termed Dset) and highlight that XConTest provides more biologically interpretable results.

## Results

### Overview of Convergence Analysis and Proposed Methods

Recent literature uses the word *convergence* loosely to refer to a marked similarity between elements under comparison. We reserve the term to mean the similarity between the profiles of gene expression after cells underwent different perturbations, i.e., convergent perturbations modify a subset of common genes (Figure 1a).

**Figure 1.**
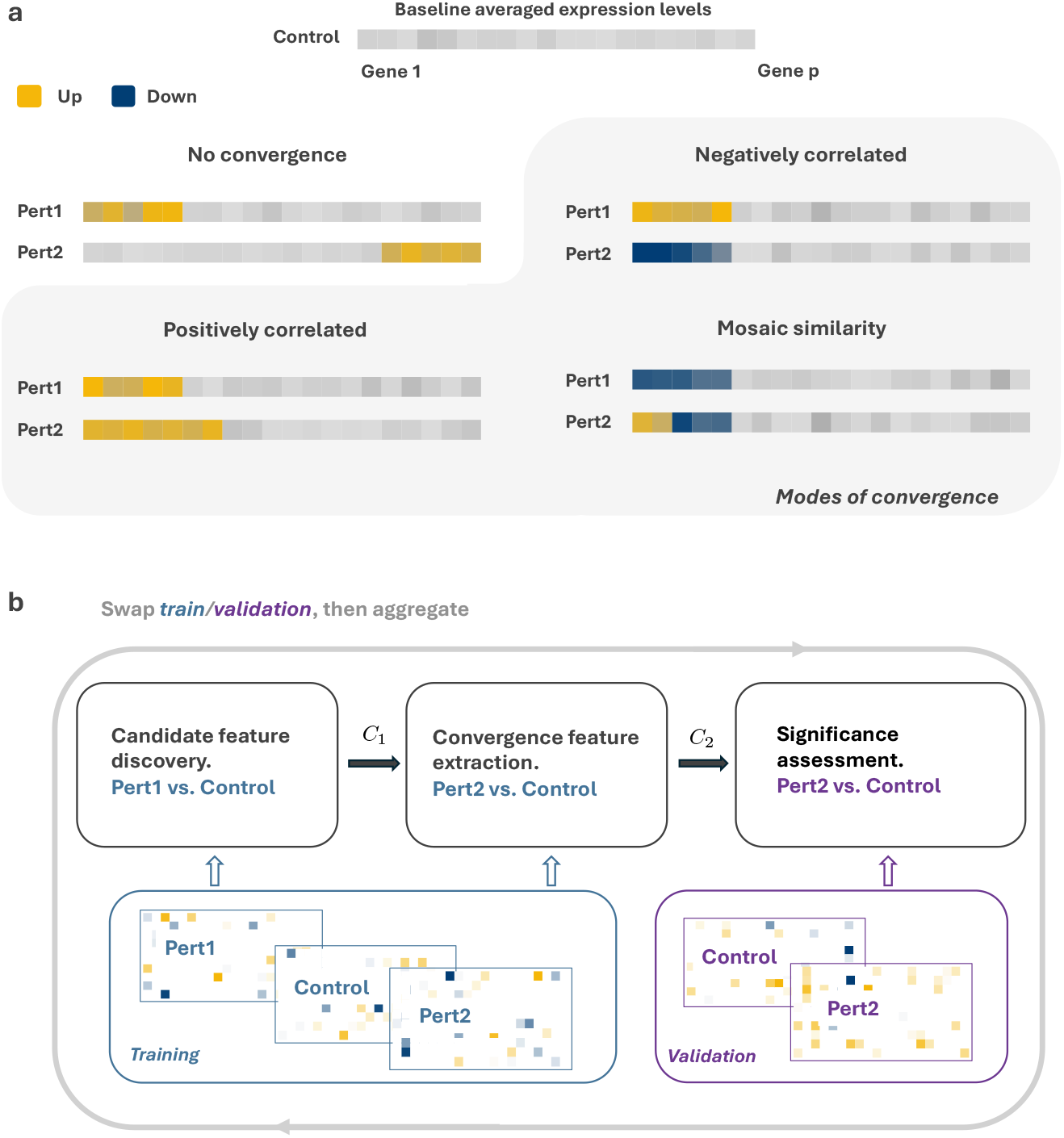
Genetic convergence and the XConTest pipeline. **a** Modes of convergence. We present three patterns of convergence as well as one example where perturbations lead to DE but not convergence. **b** A diagram of the XConTest pipeline. We use perturbation 1 to discover convergence features (gene or gene modules) and perturbation 2 to validate.

We model the cell expression profile as a high-dimensional random vector, and cells under different perturbations have different average expression levels. Let ***µ***_*c*_ = (*µ*_*c*1_, …, *µ*_*cp*_) denote the average expression levels of *p* genes for the non-targeting/control group. Suppose that there are two groups of cells that received Perturbation 1 and 2, respectively. We use ***µ***_*K*_ = (*µ*_*K*1_, …, *µ*_*Kp*_) to denote their average profiles *K* = 1, 2. Differential expression analysis examines whether ***µ***_*c*_ = ***µ***_*K*_ and if not, on which genes the difference is established.

A Convergence Analysis (CA) can be considered a second-order version of DE analysis. It attempts to answer whether perturbations 1 and 2 induce similar changes to the average expression profile, using non-targeting cells as a baseline. We use **Δ**_*K*_ = ***µ***_*K*_ − ***µ***_*c*_ to denote the change in the measured genes, CA assesses the similarity between **Δ**_1_ and **Δ**_2_. A pair of perturbations is considered convergent if: both **Δ**_1_ and **Δ**_2_ are non-zero; and there is an essential overlapping in non-zero dimensions between **Δ**_1_ and **Δ**_2_.

Convergent perturbations are believed to imply functional similarity of the perturbed genes. Figure 1a demonstrates several modes of genetic convergence. For a positively correlated pair, both perturbed genes regulate overlapping downstream pathways, and their regulation directions are identical. For the negatively correlated pair, they have identical downstream targets, whereas the regulation directions are opposite. They should still be considered convergent, and we expect a proper CA method to return positive comparison results for this case. This example also illustrates how CA is different from assessing **Δ**_1_ = **Δ**_2_. Mosaic similarity is also considered convergent, which can be seen as a generalization of the other two modes.

In this work, we introduce XConTest, a cross-fitted Convergence Test for CA problems. The method pipeline is illustrated in Figure 1b. A portion of the data—including all three groups—is used for convergent gene extraction, and some held-out samples are used to quantify the degree of similarity. Such a sample-splitting scheme can better mitigate false-discovery, and it is embedded in a cross-fitting loop to avoid sample-splitting efficiency loss. A single pass of XConTest functions as follows: (1) we collect Perturbation 1 and non-targeting cells. A classifier 𝒞_1_ is trained using the expression profiles of the *p* genes to predict whether a cell comes from the perturbed or control group. We propose using sparse-inducing linear classifiers such as Lasso [15], which can automatically select DE genes in **Δ**_**1**_; (2) another classifier 𝒞_2_ is established to distinguish Perturbation 2 and nontargeting *only using* genes selected from the **Δ**_**1**_ contrast. Typically, the classifier 𝒞_2_ uses a strict subset of genes selected in step 1, which are roughly the important DE genes for both **Δ**_**1**_ and **Δ**_**2**_; and (3) apply 𝒞_2_ to held-out data to calculate test statistics and comparison p-values.

Candidate feature extraction can be done at either the gene level or the gene module level. XConTest can integrate any algorithms that establish gene modules (programs) into its pipeline, many of which rely on empirical gene co-regulation patterns. For example, [16, 17] and [9]. In this work, we favor a pipeline including both CS-CORE [18] and WGCNA [16]. Before executing the first step of XConTest, we need to use the expression profiles to discover correlated genes and divide the full list of genes into modules. When constructing the classifiers 𝒞_1_ and 𝒞_2_, we also need to replace Lasso by Group Lasso [19] to recognize the grouping. The “features” in Figure 1 would be modules rather than individual genes.

This aggregated XConTest still controls false discovery, and the convergence inspection will be conducted at the gene-module level. We see this combination as a tool to enhance both power and interpretability. When two perturbed genes modify identical downstream modules, but do not have significant overlap at a gene level, aggregated XConTest can still detect their similarity.

Other conceptually simpler methods have also been considered when XConTest was developed. We will examine a Direct Gene Set (*Dset*) method in simulation and a real-data example as well. Dset tests whether two perturbations have overlapping DE gene modules (More technical details, including family-wise type-I error control, are presented in the Materials and Methods section).

Although conceptually simple, the Dset Method performs well in simulations when only a small number of modules are affected by the perturbations. However, in real Perturb-seq data, where many modules exhibit moderately nonzero effects, it will indicate convergence even when the main signals do not align. In contrast, XConTest better emphasizes the overall profile difference and offers easier interpretation when processing a large number of perturbations. More details will be presented in our case studies.

### XConTest controls both type-I and type-II errors

A simulation study was conducted to evaluate the statistical performance of XConTest. We use data in [20] to train a generative model that can generate expression profiles of non-targeting cells, and illustrate that the generated cells’ expression level matches the original data on a subset of 1,500 highly variable genes (Fig. 2a). A subset of the cells has a subset of genes modified to mimic perturbed groups. The empirical distribution of XConTest’s test statistics is compared to the standard normal (sample size = 200, number of genes = 1,500, Fig. 2b). When the sample size is comparable or smaller than the number of measured genes, many statistical procedures do not have easily tractable distributions —however, XConTest is designed to have an approximately Normal distribution under the null hypothesis, i.e., when there is no convergence.

**Figure 2.**
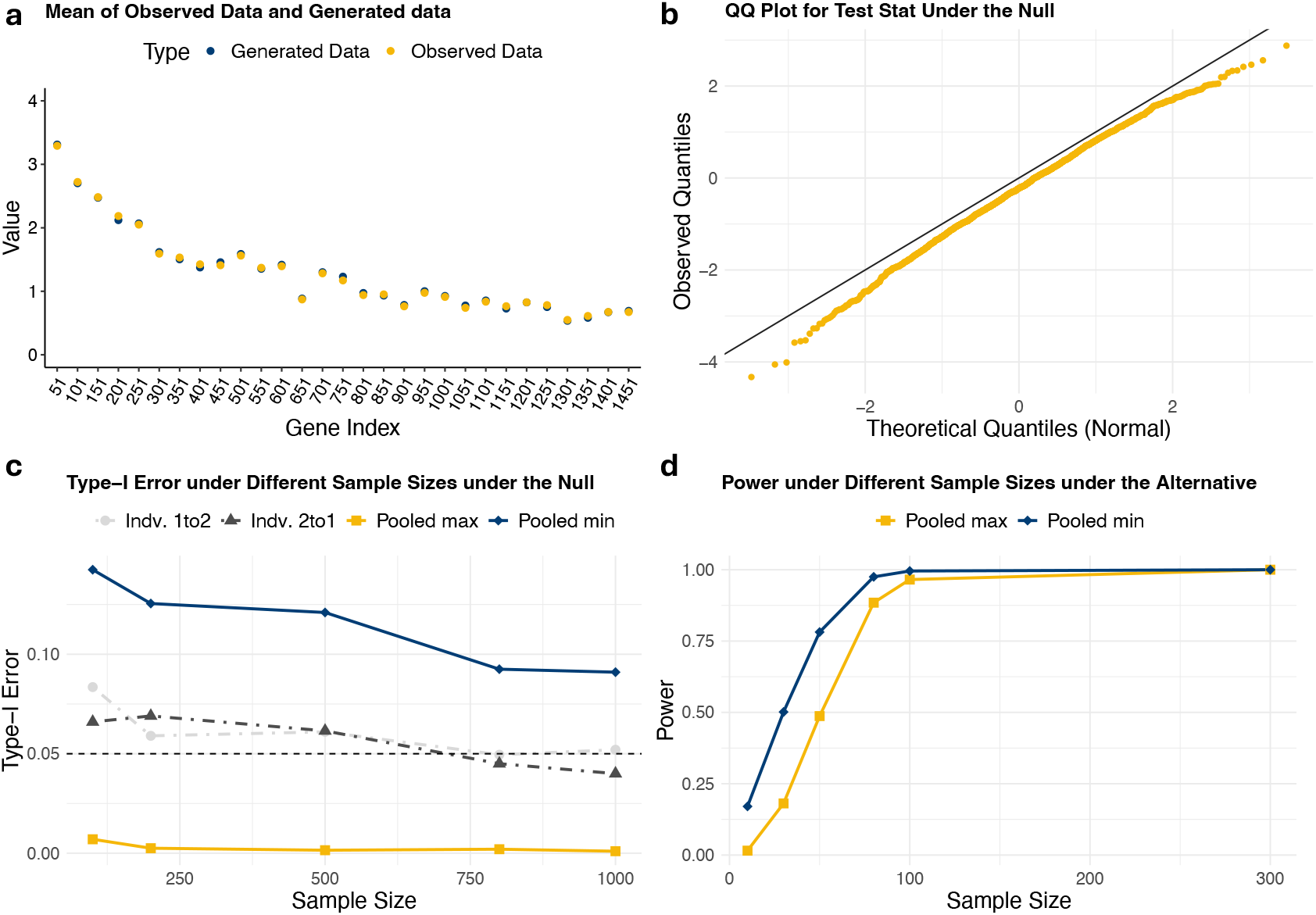
Simulation Study on XConTest. **a**. Gene mean expression of observed cells and generated cells that are sampled from a diffusion model trained. The gene expression levels overlap well. **b**. Q-Q (quantile-quantile) plot demonstrates that the test statistic is approximately Normal under the null hypothesis. **c**. Type-I and **d**. Power of XConTest under the null and alternative hypotheses. We include results from both individual and pooled p-values.

XConTest is a directional method. When data is abundant, it is insensitive to which perturbation group is used for candidate feature discovery (and which one is for the later feature extraction and significant assessment). However, in practical settings, we observe that XConTest may give different inferential results when swapping the roles of perturbation groups. This is expected when they do not induce almost identical DE patterns. We denote the direction presented in Figure 1b as 1 → 2 and reversed as 2 → 1. Figure 2c illustrates that both directions offer well-calibrated testing procedures (with a slight inflation when sample size *<* 250). One may consider either *p*_min_ = min(*p*_1to2_, *p*_2to1_) *< α* or *p*_max_ = max(*p*_1to2_, *p*_2to1_) *< α* as the significant criterion. The previous states that so long as one direction is successful, we determine the given perturbation pair is convergent; and the latter requires both directions to be predictive. In practice, we recommend using the stringent criterion based on *p*_max_ (Fig. 2c). Theoretically, *p*_min_ can control the overall type-I error less than 2*α*, but in general, it requires an unrealistically larger sample size. The maximum offers less power as expected, but it has reached one when a moderate number of cells have been collected, where the convergent genes only have 10% less expression than the baseline (Fig. 2d). The rest of this paper applies *p*_max_ *< α* when reporting significance results. More details on this simulation study are provided in Materials and Methods.

### Aggregated XConTest Permits Gene-Module Level Convergence Analysis

We conducted a simulation study to illustrate the application of the gene-module level XConTest. Specifically, we generated expression profiles of 1,500 genes for six treatment groups and one control group. These genes were grouped into 30 gene modules—we are using a simple sequential grouping, but in practice, this step should be done in a data-driven fashion and factor in gene-gene correlation—among which around 20% have modified expression level for each perturbed group (labeled A–F in Fig. 3a). Each of the perturbed groups has 200 cells, and the control has 400.

**Figure 3.**
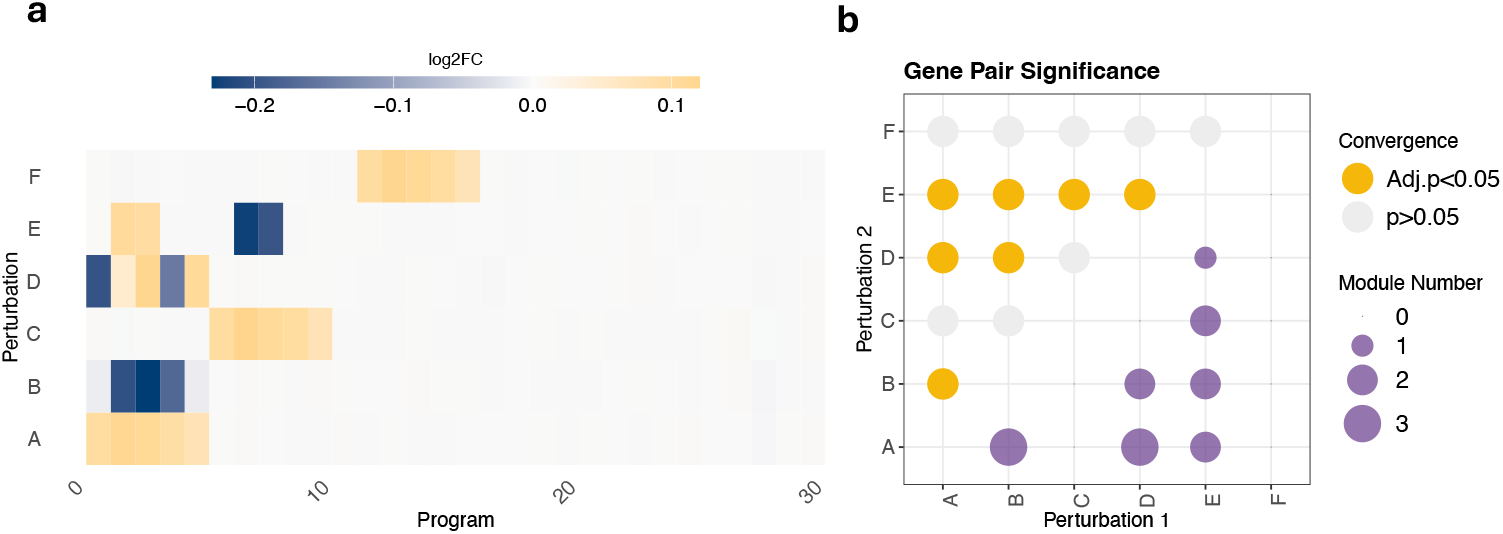
Results of aggregated XConTest on simulation data with perturbed gene modules. For each gene module, 20% of genes are perturbed and the rest are set unchanged as the non-targeting group. Artificial Perturbation by detecting gene-module convergence. **a**. shows the heatmap of the average logarithm fold change of gene expression level in each module. **b**. shows the testing results by XConTest, where all true signals and no false signals are discovered.

The heatmap in Figure 3a is a common visualization in the literature for perturbation convergence (e.g., [9]). The color intensity represents the average log fold change of gene expression within each module (average over samples, then over genes in each module).

We applied both the aggregated XConTest and Dset to the simulated datasets. The convergence results and the number of Frequently Selected Features (FSFs)—features selected in *>* 50% cross-fitting fold— are recorded (XConTest in Figure 3b, upper and lower triangular, respectively, and Dset in Figure S1). For the simple XConTest, each feature is one single gene, whereas for the aggregated XConTest it is one gene module. Recall that for the aggregated XConTest, coupled with group Lasso, we select either none or all of the genes in the “convergence feature extraction” step to establish 𝒞_2_ (Fig. 1). Although the fold change is less than 20%, this simulation setting is still not challenging, and both of the methods give similarly good recovery. Both of them offer valid type-I error control—for example, F is not convergent to any other perturbations. Pairs establishing signals are also successfully identified.

A subtle yet important distinction between XConTest and Dset emerges when comparing Perturbations A and B. In our simulation setup, all of the first five modules were modified for perturbation B. However, modules 1 and 5 carry only weak signals—which appear to be faint blue in Figure 3a. The two methods report different numbers of FSFs for the A–B comparison. Dset detects all five modules (1–5) as convergent features, illustrating its strong testing power. In contrast, XConTest reports only the three modules with substantial signal as its FSFs. This is expected as its sparsity-inducing classifiers 𝒞_1_, 𝒞_2_ exclude modules 1 and 5 for providing little additional predictive capacity once modules 2–4 are included. XConTest avoided reporting too many convergent pairs when such weak signals are more ubiquitous in real data. This difference is further demonstrated in our Perturb-seq example.

### Application: CRISPRi Data from a Neuronal Differentiation Study

There has been tremendous progress in identifying Autistic Spectrum Disorders (ASD)-associated genes. However, an open question remains whether the identified genes induce unique transcriptional signatures or whether their mechanistic impacts are similar and comparable. In [21], the authors coupled CRISPR-Cas9 transcriptional repression with single-cell RNA sequencing to study the pathological mechanisms of 13 ASD-related genes—most of the repressed genes have more than three published studies supporting their implication. Single-cell transcriptomic profiles were collected at a series of neural differentiation time points. We applied gene-level XConTest to assess perturbation similarity using data from the late pseudotime stage, when the cell differentiation is more mature.

The results of the pairwise convergence analysis of the selected perturbations are presented in two ways (Fig. 4a): binary significance results (upper triangular); and the number of genes overlapping (i.e., the FSFs from XConTest for each pair; lower triangular). We conducted a Gene Ontology(GO) analysis of the FSFs for the ADNP-PTEN pair to assess their functional concentration, and many of them concentrate on neural development-related pathways, revealing the functional similarity of these two genes (Fig. 4b).

**Figure 4.**
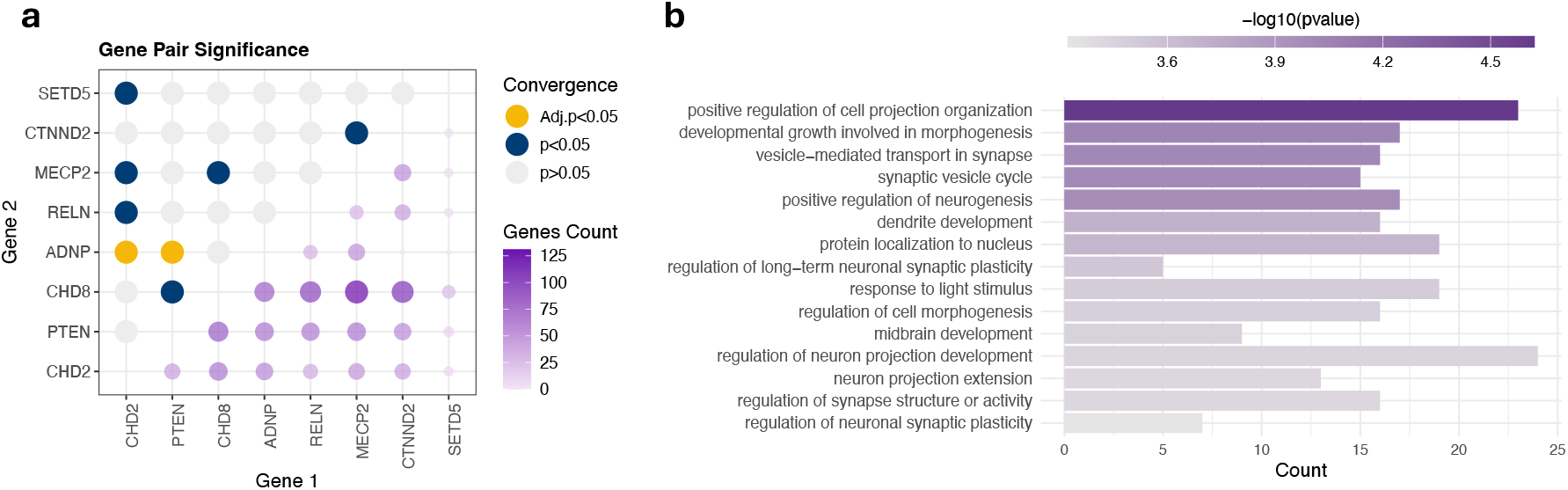
Results of Convergence Testing by XConTest on the dataset of [21]. **a**. shows the binary CA results and number of FSFs for each selected pair **b**. shows the GO analysis for genes detected where ADNP and PTEN converge.

### Application: Perturb-seq Profiling of a Human Macrophage Cell Line

Aggregated XConTest was applied to the data reported in [20] to investigate the functional similarities of 598 genes involved in the immune response to bacterial lipopolysaccharide. Datasets with such a large number of perturbations focused on a narrowly defined biological process are uncommon in the literature. In their study, the authors introduced novel experimental and computational schemes to improve the efficiency of perturbation screening and benchmarked these against standard Perturb-seq. In this investigation, we used the datasets derived from the standard Perturb-seq experiments for simplicity.

The top 50 perturbed genes were selected in [20], based on their average magnitude of effect on all downstream genes. The genetic convergence results, based on a 2k subset of the total 17k genes, are examined using XConTest (Fig. 5a). Among the 1225 (50 *×* 49*/*2) unique perturbation pairs, 927 of them do not yield significant evidence of convergence, when controlling for the type-I error at 0.05. Among the remaining pairs, 119 have a strong convergence signal (significant in Bonferroni corrected test).

**Figure 5.**
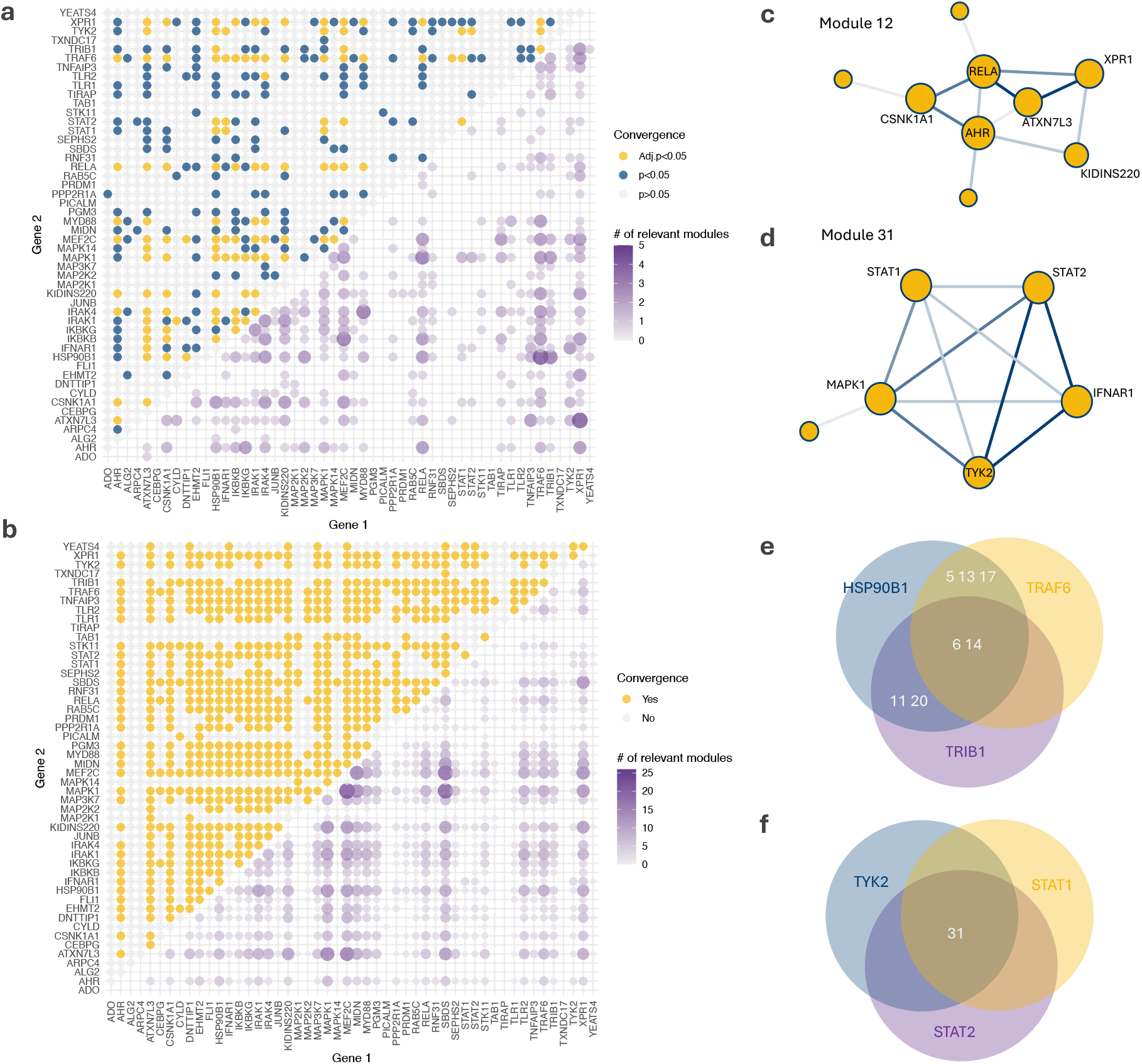
Convergence analysis of the Perturb-seq experiment reported in [20]. **a**. Convergence results of aggregated XConTest; **b**. convergence results of Dset; **(c–d)** selected gene modules and perturbations that converge on them; **(e–f)** selected convergent perturbation groups and the modules they converge on. In **e**, both modules 6 and 14 are selected by all three convergence pairs, but 11 and 20 are unique to the HSP90B1-TRIB1 pair (they converge on 4 modules in total). GO terms of modules frequently appearing in this figure are listed in Fig. 1.

As a comparison, we include the results of Dset in Fig. 5b. Recall that this method declares convergence when two perturbations have at least one differentially expressed gene module that overlaps. For these data, it claims ubiquitous convergence: 669 out of 1225 pairs are identified. Generally speaking, Dset offers more discovery opportunities— particularly valuable for data with low signal strength. However, its advantage is less evident for these conventional Perturb-seq datasets, which tend to fall within the sweet spot of XConTest. In this regime, XConTest better prioritizes perturbation pairs with stronger and more consistent evidence of convergence.

There are multiple dimensions to further examine the convergence results. Next, we stratify the convergence pairs by active modules (Fig. 5c and d). For example, an edge between STAT1 and MAPK1 in panel d indicates that: i) these two perturbations are convergent at 0.05*/*1225 type-I error level; and ii) module 31 has been selected as one of the FSFs in both “STAT1 to MAPK1” and “MAPK1 to STAT1” directions, which implies a consistent signal.

We also identified clusters of perturbations that converge on multiple modules (Figs. 5e and f and S2), and several of these modules are associated with GO terms relevant to immune-related responses (Table 1). Specifically, three perturbation groups, HSP90B1, TRAF6 and TRIB1, converge on modules 6 and 14. In addition, HSP90B1 and TRAF6 converge on modules 5, 13 and 17 (so this pair converges on 5 modules); and HSP90B1 and TRIB1 also converge on modules 11 and 20. Some of the convergent pairs discovered by XConTest have been explored in the literature as well. For example, [22] provides evidence that Heat Shock Protein 90 (HSP90) contributes to bone destruction in rheumatoid arthritis by promoting osteoclast differentiation through the TRAF6 signaling pathway.

**Table 1:** Representative GO terms associated with selected modules. For each module, we report the biological pathways’ name with GO ID.

| Module Index | Representative Gene Ontology Terms |
| --- | --- |
| 6 | Inflammatory response (0006954); Cytokine production (0001816); Adaptive immune response (0002250) |
| 12 | T cell activation (0042110); Lymphocyte activation (0046649); Antigen receptor-mediated signaling pathway (0050851); T cell receptor signaling pathway (0050852) |
| 14 | Blood vessel development (0001568); Skeletal system development (0001501); Muscle cell differentiation (0042692) |
| 31 | Response to type I interferon (0034340); Defense response to virus (0051607); Interferon-mediated signaling pathway (0140888); Cytokine production (0001816) |

Perturbation pairs that consistently converge on a small number of modules also highlight meaningful functional similarity. STAT1, STAT2, and TYK2 are part of the JAK/STAT signaling pathway [23, 24]. TYK2, one of the four JAK kinases, participates in phosphorylating the membrane cytokine-receptor upon ligand binding. STAT1 and STAT2 then dock at these sites, are phosphorylated by JAKs, and activate further downstream gene expression. All three genes are well known for their central roles in the type I interferon signaling pathway [25, 26, 27]. The convergence results based on current Perturb-seq data support this functional similarity: all three gene pairs show convergence, and only the module (31) that is associated with type I interferon has been consistently selected. When examining large-scale convergence results, both broadly convergent pairs and highly specific convergences invite close attention, as each may point toward important biological mechanisms and opportunities for discovery.

### Connecting XConTest Findings with Phenotype and Pathway Information

To obtain a functional characterization of genes it would be ideal to link genetic convergence with phenotypic outcomes, especially outcomes linked to disease. [20] identified perturbations that are associated with particular phenotypes using scLinker [28]. The approach involves mapping published GWAS signals for a phenotype to genes, and then examining the signal observed at genes affected by each perturbation. Based on this mapping, the strength of the accumulated GWAS signal is assessed (Table 4b, [20]). We focus on the downstream genes that exhibit a decrease in expression due to the knockout perturbation; these are called positive results because the outcome is positively associated with the perturbed gene. We use this data source to lend interpretation to our convergence findings. We investigated results for five phenotypes strongly associated with immune-related perturbations: eosinophil percentage, Inflammatory Bowel Disease (IBD), eczema, rheumatoid arthritis, and primary biliary cirrhosis (Figs. 6a and S3).

**Figure 6.**
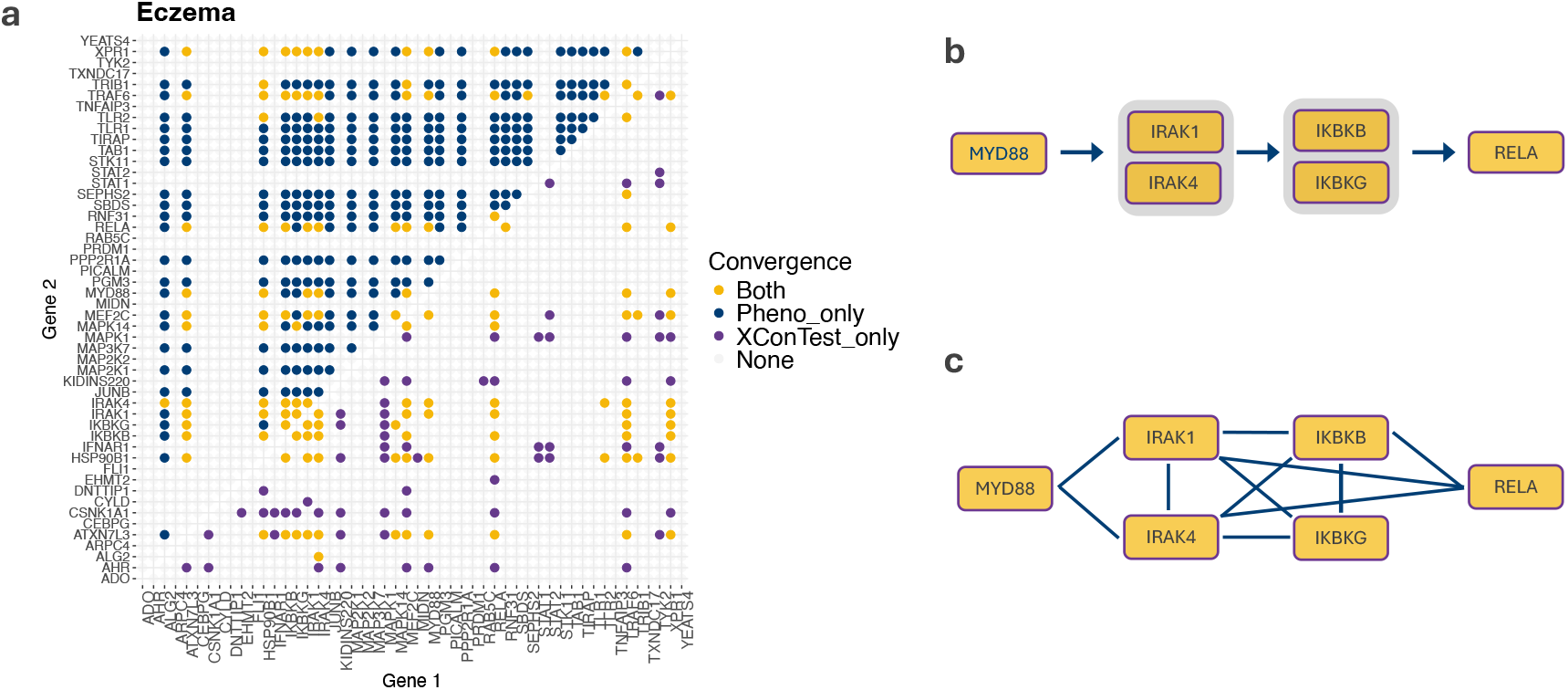
Compare XConTest findings with exogenous information. **a** Comparison of convergence discoveries by scLinker (upper triangle) and both XConTest and scLinker (lower triangle). **b** Regulatory ordering of perturbations included in this study within the MYD88-dependent NF-*κ*B pathway. **c** Joint convergence based on scLinker and XConTest closely resembles the mechanism described in **b**.

The perturbations HSP90B1, TRAF6, and TRIB1 were shown to converge by applying XConTest (Fig. 5e). Moreover, each of them can also be related to the eosinophil percentage and Rheumatoid Arthritis (RA) via scLinker as well (Fig. S3). Eosinophils are classic effector cells in inflammation, especially in allergic and parasitic contexts. They release cytotoxic granules and lipid mediators that drive inflammation [29]. Eosinophils actively produce and secrete cytokines such as IL-4,5,13, GM-CSF, and TNF, contributing to modulation of immune responses [30, 31]. While eosinophils are innate immune cells, they influence adaptive immunity. They present antigens in some contexts, produce cytokines that shape T helper (T_h_) cell differentiation (especially T_h_2), and regulate B-cell responses [32].

RA becomes chronic when the adaptive immune response fails to maintain self-tolerance and perpetuates synovial inflammation. Autoreactive T and B cells often amplify this process and drive autoantibody production. Later, elevated levels of TNF-*α*, IL-1, and IL-6 sustain a cytokine feedback loop that recruits additional immune cells and promotes joint damage [33, 34].

XConTest selected module 6 as one of the FSFs for these three perturbations (HSP90B1, TRAF6, and TRIB1), whose GO enrichments have a similar interpretation as the functions of eosinophils and pathophysiology of RA (Table 1).

Perturbations STAT1, STAT2, and TYK2 were shown to converge on module 31 (Fig. 5d), which has the GO enrichment shown in Table 1. scLinker indicates STAT1 is associated with IBD and IBD is strongly associated with all four processes identified with module 31—type I interferon response, antiviral defense, interferon-mediated signaling, and cytokine production—through immune dysregulation and inflammatory signaling in the gut [35, 36, 37, 38]. Our convergence results suggest that STAT2 and TYK2 may also play a role in IBD via JAK/STAT signaling pathway and the type I interferon signaling pathway. Indeed, the GWAS signal for both of these perturbations, as assessed by scLinker, is borderline significant.

XConTest also identified convergent patterns that aligned well with known cellular regulation pathways. In particular, many of the top convergent pairs mapped to the widely studied MyD88–dependent signaling cascade [39, 40, 41], which relates upstream receptor activation to the NF-*κ*B and MAPK transcriptional complexes. We illustrate a simplified ordering of the key regulatory components in Figure 6b, including multiple genes relevant to the current study (a comprehensive pathway diagram can be found in [40]). XConTest produced a tightly connected convergence graph for these genes (Fig. 5a), where edges represent statistically significant convergence. Only a small number of perturbations outside the known pathway exhibited consistent convergence with any of the illustrated perturbations. In contrast, Dset showed no such specificity (Fig. 5b).

In summary, when a set of convergent perturbations also shares association with a phenotype or a known regulatory pathway, it lends support to the conjecture that convergent genes have common functionality. For example, several of the genes in the MyD88 pathway are implicated in eczema by scLinker (Fig. 6a). At the same time, ATXN7L3, a gene that is not known to be part of the MyD88 pathway, converges with many of the genes in the pathway and is also associated with eczema. By providing insight into how ATXN7L3 is associated with eczema, the XConTest analyses may suggest a potential indirect role of the gene in the MYD88 pathway.

Finally, we examine parallel analyzes using Dset (Fig. S4). Although Dset identified a large number of convergent perturbations—many of which may represent weak associations—scLinker still detected additional pairs not captured by Dset. This can happen when a pair of perturbations has distinct genetic signals associated with their targets, but their targets do not overlap. Conversely, some pairs were identified by Dset and not scLinker. A notable example is MAPK1: Dset classified this perturbation as convergent with nearly all others, while scLinker did not identify MAPK1 for any pair of perturbations. Using XConTest, MAPK1 also appeared convergent with several perturbations. Although MAPK1 is not known to be directly linked to the phenotypes examined here, it encodes a core component of the mitogen-activated protein kinase (MAPK) pathway, which interacts with the JAK/STAT signaling cascade [42], suggesting potential indirect associations. The absence of MAPK1-related convergence in scLinker could imply that genetic variation within MAPK pathway genes does not contribute to the phenotypes under study. Alternatively, MAPK1 influences these traits indirectly, as evidenced by its convergence with key genes (e.g., STAT1). However, downstream MAPK1 targets are more evolutionarily conserved, and therefore their expression is less likely to be governed by regulatory genetic variation. Consistent with the latter hypothesis, MAPK1’s targets exhibit low LOEUF scores [43], providing at least partial support for this interpretation (Fig. S5, 2-sample t-test comparison of means, *p* = 10^−4^). Together, these results suggest that identifying convergent perturbations can complement GWAS in elucidating gene function and highlight avenues for future investigation.

## Discussion

Understanding the functional characterization of genes in human cells has long been a central goal of genomics and systems biology, yet the function of a substantial proportion of genes remains poorly understood [44]. Recent advances in pooled CRISPR-based perturbation screens with molecular readouts now enable systematic inference of genetically causal networks. By integrating and iteratively refining insights from such perturbation studies, it is becoming increasingly feasible to delineate causal relationships among genes at scale [45]. With XConTest, we address a key aspect of this endeavor—the identification of CRISPR perturbations that exert convergent effects on gene regulation, that is, perturbations influencing a shared set of downstream targets. Unlike heuristic strategies, XConTest provides a rigorous statistical framework to quantify the convergence of gene functions.

We considered two complementary approaches for identifying convergent perturbations. Dset provides a sensitive means of detecting shared functional outcomes, even when the magnitudes of perturbation effects differ substantially. In contrast, our framework employs a sparsity-inducing group Lasso formulation that explicitly penalizes weak effects across modules. This structure enforces cross-module comparability, such that only perturbations with consistently strong signals are propagated to subsequent inference steps. Consequently, convergence is declared only when strong effects align across perturbations, yielding a conservative yet statistically rigorous measure of functional similarity.

GWAS have identified thousands of human genetic variants associated with various diseases and traits, yet the majority of these variants reside in noncoding regions with unknown target genes and functions. CRISPR-based perturbation experiments now provide a powerful means to interrogate the functional consequences of such variants [46]. [9] introduced an innovative approach that links genetic variants to their putative target genes and leverages Perturb-seq to identify convergent transcriptional effects across perturbations—and, by extension, across variants. XConTest builds on this direction by offering an inferential framework to quantify functional convergence among perturbations. Nevertheless, additional work will be needed to establish an integrated, scalable pipeline for translating these experimental and computational advances into practical applications.

In this study, we focus our analysis on perturbations with scRNA-seq readouts; however, emerging technologies now enable a wide range of multimodal perturbation assays. These include chromatin-based readouts with Perturb-ATAC [47], protein-level profiling with ProCODE [48], joint RNA–protein measurements with ECCITE-seq [49], CaRPool-seq or Perturb-CITE-seq [5], as well as combined RNA–chromatin profiling with Perturb-SHARE-seq [51]. XConTest is readily extensible to these settings: it can be applied to any high-dimensional or longitudinal outcome data, provided that baseline (control) and perturbation (treatment) groups are clearly defined. Generative models have been developed to predict the outcome of perturbation experiments that have not been performed [52, 53, 54]. [45] envision these pseudo-data as one step toward producing a perturbation cell and tissue atlas. Part of attaining that goal would involve including these pseudo-data in the analysis pipeline of XConTest. This merits further investigation.

## Materials and Methods

### Data Acquisition

We evaluated our methods using two genome-scale CRISPR datasets. The Neuronal Differentiation Study targets 14 neurodevelopmental genes, including 13 autism risk genes, in LUHMES human neural progenitor cells [21]. After CRISPR targeting, cells were differentiated into postmitotic neurons and their transcriptomes were profiled using droplet-based scRNA-seq [55]. The data related to this work was downloaded from NCBI Gene Expression Omnibus (GEO; https://www.ncbi.nlm.nih.gov/geo/) under accession number GSE142078. We implemented the preprocessing scripts provided in the supplementary materials of [21].

The Human Macrophage Cell Line dataset assesses the impact of 598 genes in the immune response to bacterial lipopolysaccharide [20]. The study investigated the performance of two experimental knock-out procedures, but we restricted our analysis to data derived from conventional perturb-seq methods. The related data was downloaded from GEO GSE221321.

### The XConTest Method

XConTest partitions the data into multiple folds, using a subset of the samples to identify potentially convergent genes and reserving the remaining samples to quantify the signal strength. In practice, the number of folds, denoted by `NumFold`, is typically set between 5 and 10 to balance estimation quality and computational efficiency.

Below, we provide further details on the proposed XConTest framework.

1. We collect transcriptomic profile data from the control, perturbation 1, and perturbation 2 groups, withholding a (`NumFold`)^−1^ proportion of samples from each group for subsequent significance assessment. This held-out data is referred to as the validation data.
2. In the training step, we have three collections of single-cell profiles: 𝒟_p1_, 𝒟_p2_, and 𝒟_con_, corresponding to the Perturbation 1, Perturbation 2, and Control groups, respectively. Each cell is represented by an expression vector ***X*** = (*X*_1_, …, *X*_*p*_)^⊤^ ∈ ℝ^*p*^, where *p* denotes the number of genes. When pooling any two of these datasets, the combined samples can be treated as random draws from a mixture distribution of two populations.
3. We append a categorical group label *Y* ∈ {`con, p1, p2`} to each ***X***. We first fit a logistic Lasso model regressing *Y* on ***X*** using 𝒟_con_ and 𝒟_p1_:

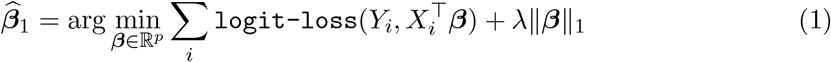

where the summation is over all samples *i* ∈ 𝒟_con_ ∪ 𝒟_p1_. The sparse regression coefficients 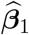 implies the classifier *C*_1_ in Figure 1b. We denote the support of 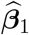, that is, the set of genes with non-zero estimated coefficients, by *s*_1_ ⊂ 1, …, *p*.
4. Fit another logistic Lasso model, regressing *Y* on ***X*** using 𝒟_con_ and 𝒟_p2_. The coefficients are denoted as 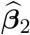 and its formula is almost identical to that of 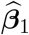: but it is restricted to have zero in all dimensions outside *s*_1_. The coefficients 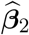 defines the classifier 𝒞_2_ in Figure 1b. Compute the leading Principal Component (PC1), ***υ̂***, using 𝒟_con_. Similarly, we only allow genes in *s*_1_ to have non-zero loadings. In practice, the constraints on 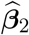 and ***υ̂*** are readily implemented by trimming dimensions outside *s*_1_, applying standard software for logistic regression or PCA, and padding zeros to recover the full *p*-dimensional vectors.
5. Define the CA projection direction as (normalized to have a unit Euclidean norm):

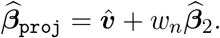

where *w*_*n*_ is a weighting parameter that diverges with increasing sample size. In this study, we set *w*_*n*_ = *n*^1*/*3^, where *n* denotes the smallest sample size among 𝒟_p1_, 𝒟_p2_, and 𝒟_con_.
6. Let 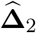 denote the estimated sample mean difference between the perturbation 2 and the control groups. We project this estimated difference onto 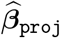 to obtain

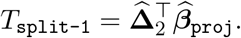

The corresponding standard error is estimated as

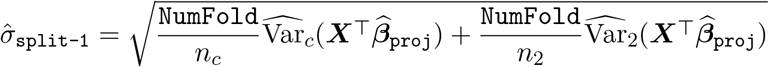

where *n*_*c*_, *n*_2_ are the total sample sizes from control and perturbation 2. The operator 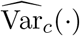 represents the sample variance of the projection scores computed using control cells in the validation set, and 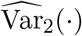 is defined analogously for the perturbation 2 cells.
7. Finally, we repeat steps (1)–(6) across all cross-validation folds to obtain fold-specific test statistics *T*_split-*j*_, *j* = 1, …, NumFold. The overall test statistic is computed as

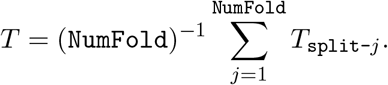

Its standard error is estimated by

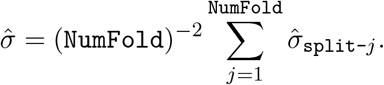

The resulting studentized statistic, 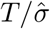 is approximately standard normal under mild regularity conditions. A formal theoretical justification is provided in Corollary 3.2 of [56]. Notably, while many high-dimensional statistics deviate from normality asymptotically, the test statistic in our framework retains a Gaussian limiting distribution.

### Aggregated XConTest

We recommend conducting CA on the gene-module level as demonstrated with the data from [20]. The pipeline is very similar to that of gene-wise XConTest. The aggregated XConTest has one extra preprocessing step:

(pre) Divide the *p* genes into gene-modules using any available methods. We implemented the integrated including both CS-CORE and WGCNA (a vignette is available at https://changsubiostats.github.io/CS-CORE/articles/CSCORE.html). For genes not strongly associated with any others, we further divide them into smaller modules based on their GO similarity.

The only modified steps are (3) and (4). Instead of fitting logistic Lasso, we need to incorporate the grouping information and solve the group Lasso optimization:

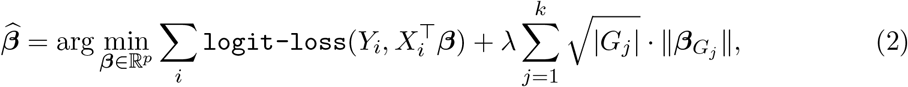

where *k* is the number of gene modules, |*G*_*j*_| is the number of genes in the *j*-th module and 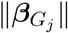 is the norm of ***β*** restricted on group *j* only. We applied the software in R package `grpreg` [57]. The set *s*_1_ in step (3) are all the genes in the selected modules of 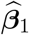.

### The Dset method

In this section, we describe the details of the Dset method that we used as a comparison to XConTest in several examples. Pairwise Dset comparison consists three steps

1. Divide the full list of genes into modules. We will implement CS-SCORE in this step as well, for the ease of comparison to XConTest. This step would be idetical to the (pre) step of aggregated XConTest.
2. Apply a DE test to each of the established modules. In this work, we considered the powerful test introduced in [58].
3. If both Perturbation 1 and 2 establish a significant DE signal on the same module, we claim that they are convergent. The significance level needs to be carefully adjusted to control familywise type-I error, detailed below.

Let *N* denote the number of perturbations under comparison. And let *M* be the number of gene modules established in step (1).

Prob(Dset reports at least one pair converge | *H*_0_)

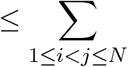 Prob(Dset reports perturbations *i* and *j* converge | *H*_0_)

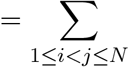 Prob(At least one module reported to be DE for both *i* and *j* | *H*_0_)

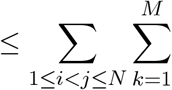 Prob(*i* and *j* are both DE on module *k* | *H*_0_)

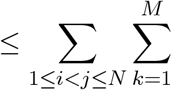 Prob(*i* is DE on module *k* | *H*_0_)

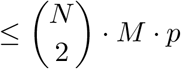

To control the family-wise type-I error at level *α*, we need to set the significance level of each DE test at no larger than 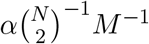. In the artificial perturbation setting (Figure 3), *M* = 30, *N* = 6.

### Simulating Data via Diffusion models

Diffusion models, as an important type of generative model, are trained on data from a target data distribution that we aim to approximate or sample from.

Completing the generation of new samples can be divided into two stages: 1) we will gradually add some noise to the training data (forward procedure), and then 2) denoise for sampling, using the estimation of the score function, which is a crucial tool to be discussed later (backward procedure). Formally, the forward procedure can often be expressed as generating a chain of random variables *X*_1_ → *X*_2_ → · · · → *X*_*T*_ from the data distribution *p*_data_ as follows:

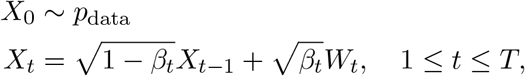

where {*W*_*t*_}_1≤*t*≤*T*_ denotes a sequence of independent noise vectors drawn from 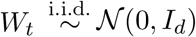, and *β*_*t*_ ∈ (0, 1). As a result, the data will be contaminated by random noise and finally approximate the standard normal distribution.

There are multiple ways to execute the backward procedure. In this work, we implement the widely-applied DDPM (Denoising Diffusion Probabilistic Model) framework proposed in [59]. This algorithm utilizes the following backward sampling procedure: starting *Y*_*T*_ ∼ 𝒩 (0, *I*_*d*_) and a user-specified *α*_*t*_ ∈ (0, 1),

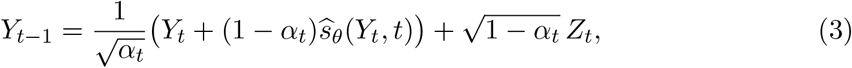

for 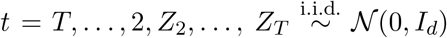. Here *ŝ*_*θ*_(·, *t*) denotes an estimator of the score function for the distribution of *X*_*t*_, defined as ∇ log *pX*_*t*_ (*x*). Equation (3) could be seen as denoising *Y*_*t*_ into *Y*_*t*−1_ gradually with the help of *ŝ*_*θ*_, an approximation of the score function. The parameter *θ* represents parameters in the neural network. DDPM ensures a good quality of sampling as long as *ŝ*_*θ*_(*Y*_*t*_, *t*) is a good estimator of the score function.

Vanilla diffusion models can only sample from a homogeneous population. For example, if the data distribution *p*_data_ contains cells under both the non-targeting group and the treatment group, the model may struggle when the user wants to generate cells of the non-targeting group only. In practice, this often requires training a separate model solely on non-targeting cells to achieve high-quality generation. A more efficient approach would allow users to specify the desired sub-category of data (cell type or perturbation condition) during the sampling procedure after training a single diffusion model on the full dataset containing all kinds of cells. This leads to the concept of the diffusion model for conditional sampling, where an additional variable *c* (the subcategory label) is added to the score function, changing it from ∇ log *pX*_*t*_ (*x*_*t*_) to ∇ log *pX*_*t*_ (*x*_*t*_ | *c*). We only need to allow *ŝ*_*θ*_ in to take one more argument *c* when sampling.

In practice, *c* could also be a multivariate embedding vector, instead of a numerical indicator, corresponding to each cell group. As suggested in [54], we use the PCA of the effect size matrix as the embedding for each cell type. The effect size is defined, for each gene-perturbation pair, as the logarithm of the ratio between the mean values of the treatment group and the control group, calculated from the data.

We took the mean of the embeddings of all treatment cell types as the embedding of non-targeting cells. The generated non-targeting cells have gene expressions that are very similar to those of the observed non-targeting cells (Figure 2a).

### Type-I error and power assessment

To test the approximate normality of the test statistic under the null hypothesis (no convergence), we generated cell expression profiles using a diffusion model (repeated 2000 rounds). The generated cells are divided into three groups, each containing 200 samples: a non-targeting group, a perturbation 1 group in which genes indexed 1–30 are up-regulated fivefold, and a perturbation 2 group in which genes indexed 31–60 are up-regulated fivefold. The distribution of the 2000 test statistics was examined in Figure 2b. Specifically, we illustrated the test statistics obtained by experiment 1 → 2, where the perturbation 1 group is used for candidate feature discovery and perturbation 2 for validation.

The simulation setting in Figure 2c is the same as the one in Figure 2b, where we only varied the sample sizes. When calculating the Type-I error, we accounted for the directional nature of our method and compared several approaches for pooling p-values, including *p*_min_ and *p*_max_ as defined in the main text.

The power of XConTest was evaluated under different sample sizes. To simulate convergent perturbations, we perturbed genes indexed by 1–15 by a factor of 0.95 in the perturbation 1 group, and genes 1–5 by a factor of 0.9 in the perturbation 2 group. Results are shown in Figure 2d.

### Simulation with perturbed gene modules

For this experiment, we first generated the non-targeting cell groups by sampling from a 1500-dimensional normal distribution. The entries are independent with marginal means of 10. Each value drawn from the normal distribution was rounded to the nearest integer and treated as a count. We fitted Poisson regression models to assess the log-fold change To generate cells simulating perturbations A to F, we modified different gene modules, each consisting of 30 consecutive genes. To reflect the biological assumption that a single perturbation influences only a limited number of genes, we restricted perturbations to 20% of the genes in each module. Perturbed cells were also drawn independently from the normal distribution with a different mean value, ranging from 4.5 to 15, and each value was rounded to the nearest integer. The mean value higher than the baseline shows the up-regulated perturbation effect while the one lower than the baseline shows the down-regulated perturbation effect. The convergence results of XConTest for this experiment were presented in Figure 3b.

## Supplementary Materials

**This PDF file includes** Figure S1 to Figure S5.

## Acknowledgments

We thank Bernie Devlin for his assistance in contextualizing the findings within the scientific literature,

## Author contributions

TZ and KR conceptualized the study; TZ designed the methodology and ES refined the testing procedures. TZ and ES created the visualizations; TZ developed the software; All authors contributed to the drafting of the manuscript and discussion of the obtained results. All authors read and approved the final manuscript. KR obtained the funding.

## Data and materials availability

The CROP-seq and Perturb-seq datasets used in this study are publicly available and were downloaded from the GEO, under accession numbers GSE142078 [21] and GSE221321 [20]. The GSM6858447 subset of GSE221321, labeled as “THP-1 cells, conventional KO screen”, was presented in this work (https://www-ncbi-nlm-nih-gov.proxy.library.ucsb.edu/geo/query/acc.cgi?acc=GSE221321). The R package HMC implementing the XConTest method is freely available at https://github.com/terrytianyuzhang/ HMC. The source code used in our study is deposited at (https://github.com/terrytianyuzhang/genetic_convergence_SA). Tutorial for implementing XConTest can be found at https://terrytianyuzhang.github.io/HMC/HMC_convergence.html.

## Supplementary Materials

**Figure S1:**
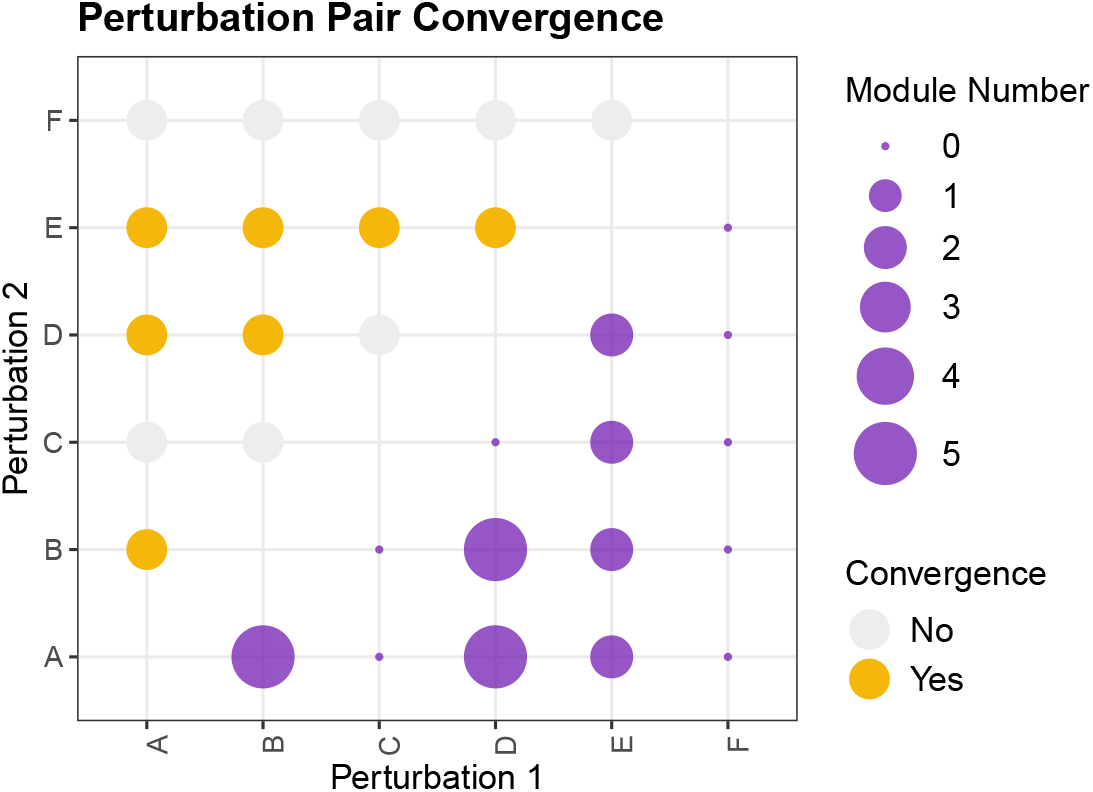
Results of Dset on simulation data with perturbed gene modules. The discovery is very to similar to that reported by XConTest in Figure 3b.

**Figure S2:**
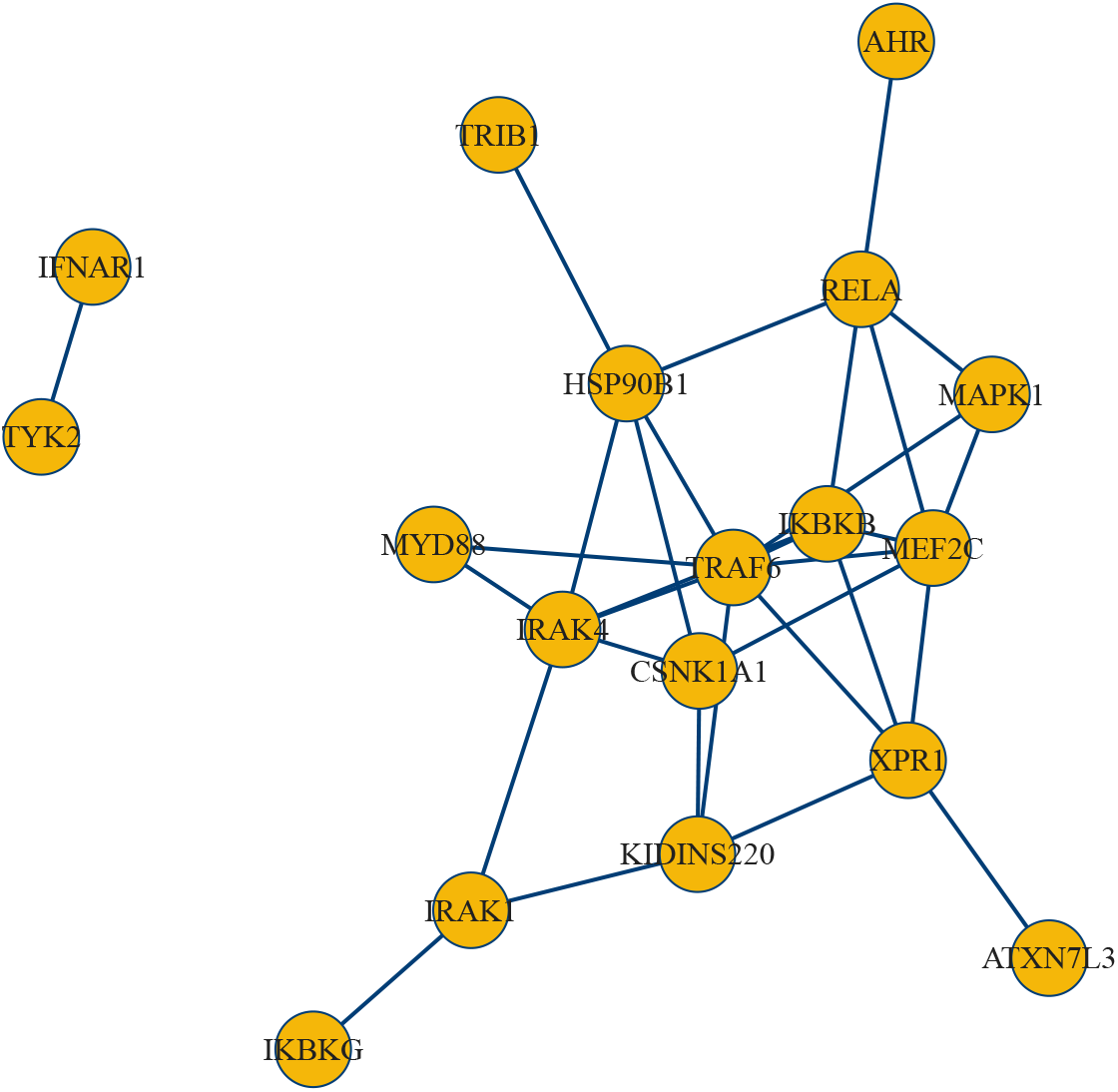
Convergent perturbations with ≥ 3 FSF-modules in the Human Macrophage Cell Line example.

**Figure S3:**
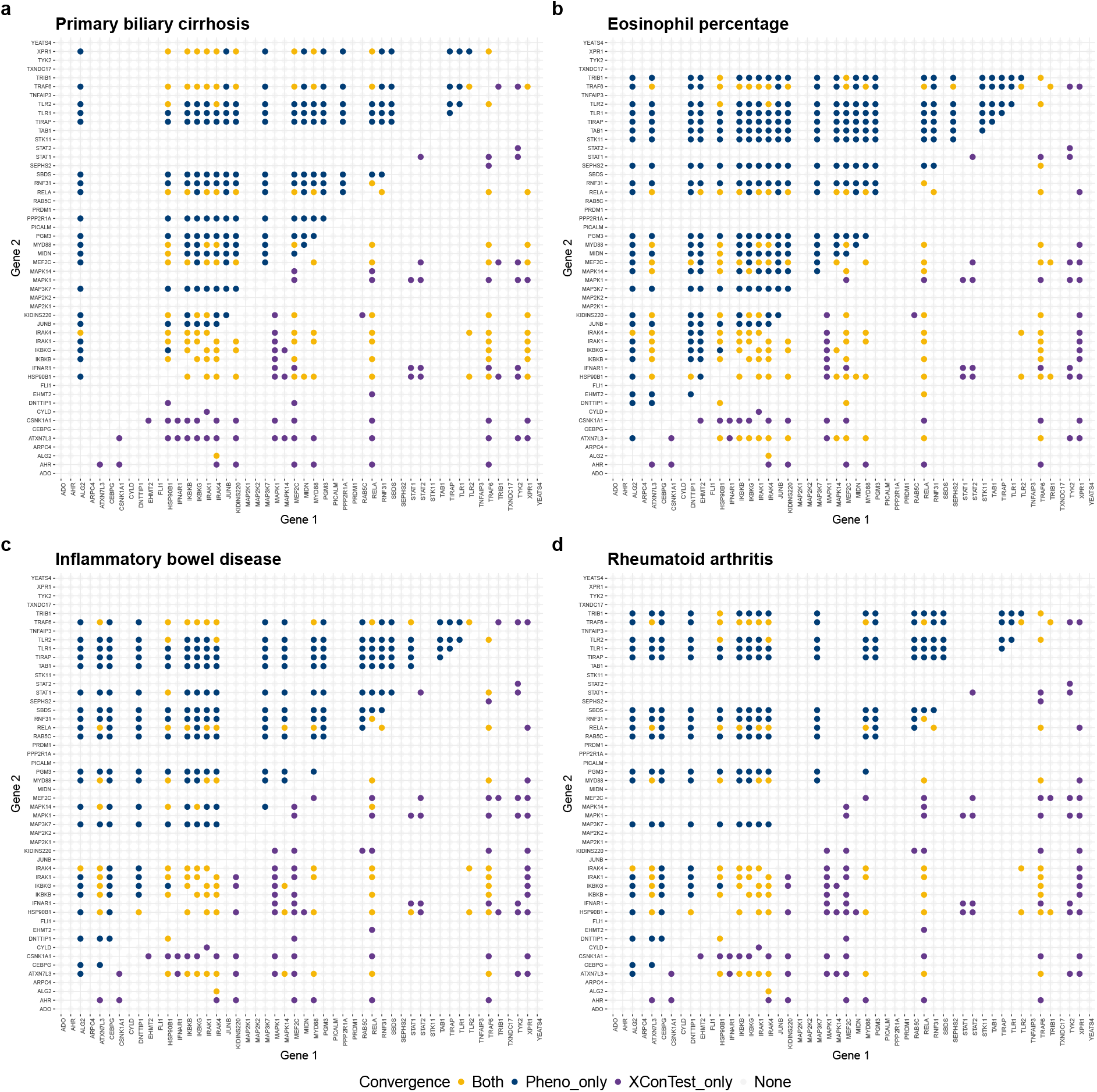
Comparison of convergence discoveries by scLinker (upper triangle) and both XConTest (lower triangle).

**Figure S4:**
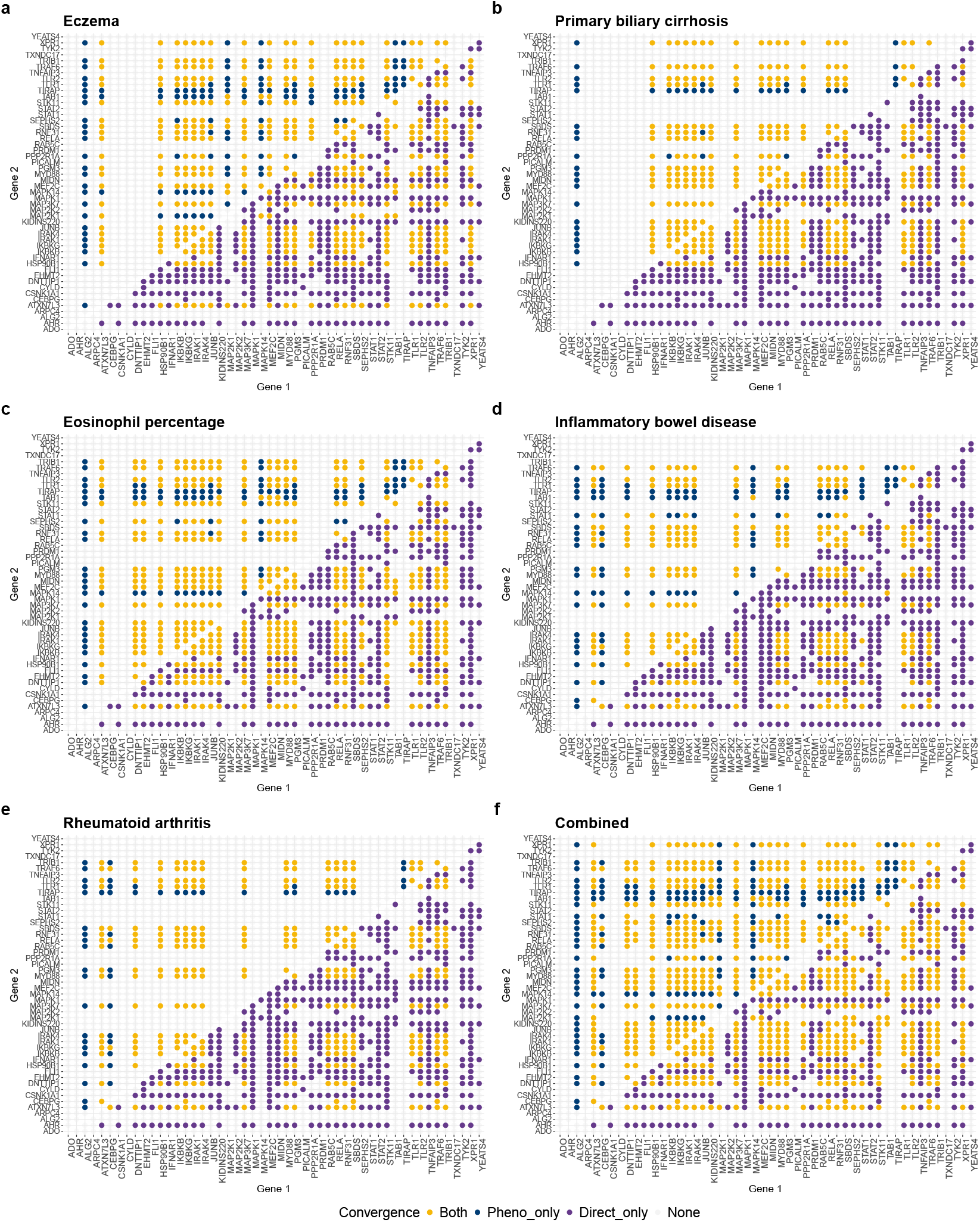
Contrast of Dset convergence with scLinker convergence for 5 phenotypes (**a**-**e**) and at least one phenotype **f**.

**Figure S5:**
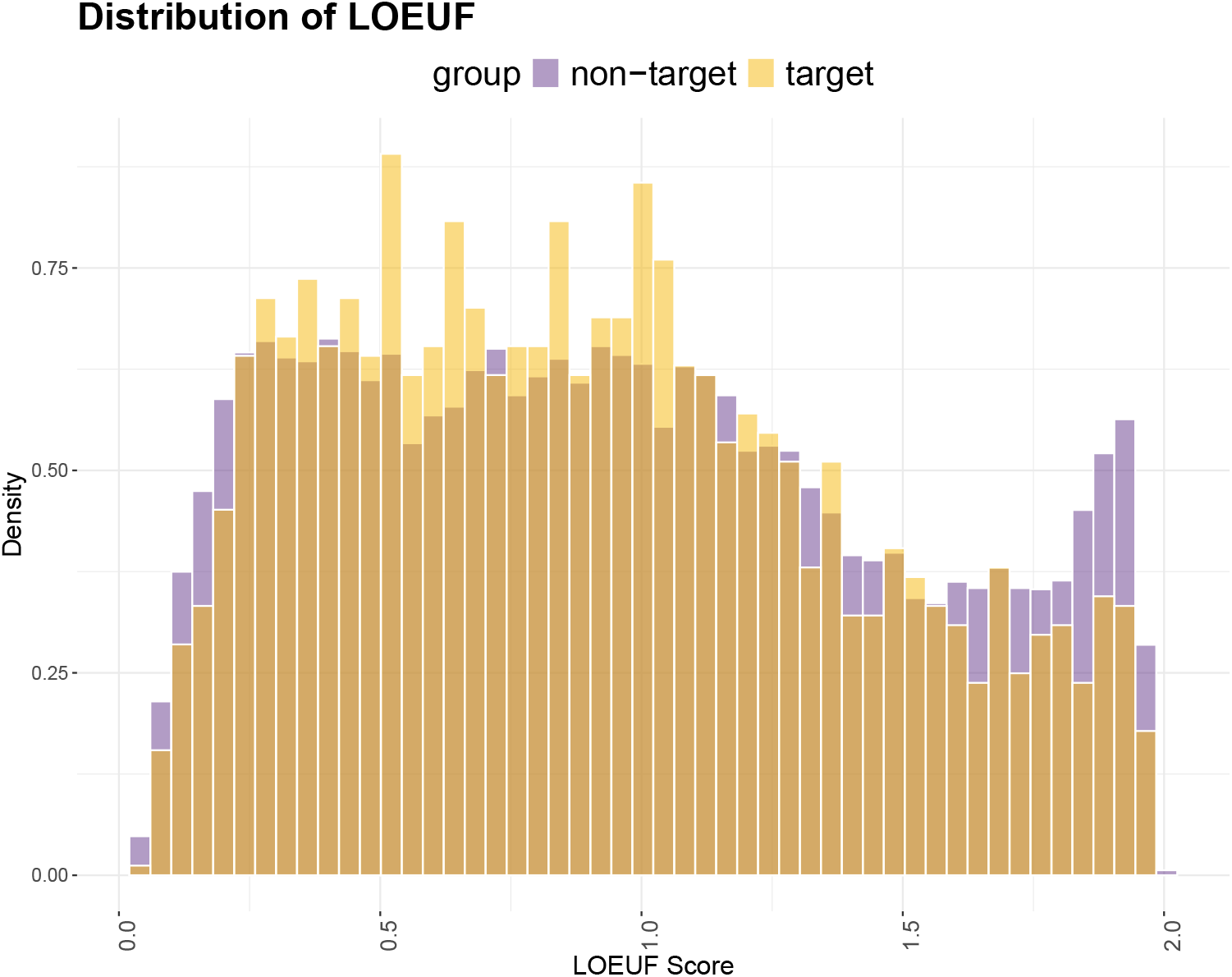
Contrast of LOEUF conservation score for target and nontarget genes of MAPK1.

